# A novel human enteroid-anaerobe co-culture system to study microbial-host interaction under physiological hypoxia

**DOI:** 10.1101/555755

**Authors:** TY Fofanova, CJ Stewart, JM Auchtung, RL Wilson, RA Britton, KJ Grande-Allen, MK Estes, JF Petrosino

## Abstract

Mechanistic investigations of host-microbe interactions in the human gut are limited by current co-culture model systems. The intestinal epithelium requires oxygen for viability, while gut bacteria are facultative or obligate anaerobes. The ability to model host-commensal interactions under dynamic oxygen conditions is critical to understanding host-pathogen interactions in the human gut. Here, we demonstrate a simple, cost-effective method for co-culturing obligate anaerobic bacteria with human intestinal enteroid monolayers under variable oxygen conditions. The Enteroid-Anaerobe Co-Culture (EACC) system is able to recapitulate the steep oxygen gradient seen *in vivo* and induce expression of hypoxia-associated phenotypes such as increased barrier integrity and expression of antimicrobial peptide genes. Using clinical strains of the commensal anaerobes *Bacteroides thetaiotaomicron* and *Blautia* sp. on established patient-derived intestinal enteroid cell lines under physiological hypoxia, the EACC system can sustain host-anaerobe interactions for at least 24 hours. Following co-culture with anaerobic bacteria, we demonstrate patient-specific differences in epithelial response, reinforcing the potential to develop a personalized medicine approach to bacteriotherapy and host-microbe interaction investigations. Our innovative EACC system provides a robust model for investigating host-microbe interactions in complex, patient-derived intestinal tissues, that facilitates study of mechanisms underlying the role of the microbiome in health and disease.

## Introduction

The human gastrointestinal tract is a site for digestion, immune system regulation, drug and nutrient absorption, and host-environment interactions ^1^. The microbiome can affect each of these processes, influencing metabolism, immune balance and therapeutic outcome ^2^. Self-organizing mini-intestines cultured *ex vivo* from intestinal biopsy samples, termed tissue-derived intestinal organoids or enteroids (used herein), present a unique opportunity for studying host-microbe interaction at the epithelial level ^3–6^. These patient-derived epithelial lineages maintain regional specificity, differentiate into all major cell types of the intestinal epithelia, reproduce physiologic activity consistent with their region of isolation, including secretion of mucus ^7–10^. Enteroids also retain the genetic background and susceptibility of the host from whom they were derived and are therefore a uniquely advantageous model for investigating the host-microbe interactions in a donor-specific manner ^11^. They also circumvent the need for animal models and follow to the principles of the “3Rs” (Replacement, Reduction and Refinement) ^12^.

Mechanistic investigations of host-microbe interactions in the human gut are severely limited by two principal challenges. First, the intestinal epithelium is oxygen dependent, while most gut bacteria are obligate anaerobes. To recreate this physiologically-relevant steep oxygen gradient across the single-cell thick epithelial layer poses a significant design challenge ^13^. Second, the intestinal epithelium is in a state of chronic low-grade inflammation due to constant exposure to luminal antigens, which are sampled by circulating immune cells ^14^. These immune cells consume residual oxygen and, coupled with the effects of counter-current oxygen exchange, keep intestinal epithelial cells in a state of constant, low-grade hypoxia known as “physioxia.” In the context of disease, such as inflammatory bowel disease (IBD), hypoxia is often exacerbated ^15,16^. Finally, the presence of microbes influences the availability of oxygen in the gastrointestinal tract; facultative anaerobes respire molecular oxygen as it becomes available whereas some commensal *Clostridia* produce butyrate that is oxidized by colonocytes to maintain hypoxia ^17^.

It has been shown that the pericellular oxygen concentration of Caco-2 cells cultured in ambient incubator conditions (18-21% oxygen) can drop to 16% oxygen when cells are cultured on membrane inserts ^18^. This high level of oxygenation observed under standard incubator conditions serves as a significant confounding factor when considering that *in vivo* oxygen levels range from 1%-11% oxygen ^13,19^. Additionally, altered gene-expression and phenotypes have been observed when comparing aerobic cultures to cultures grown at physiologic levels of oxygen, suggesting that ambient oxygen induces hyperoxic, potentially atypical, phenotypes in cell culture ^20^. Thus, being able to model host-commensal interactions under dynamic, biologically relevant pericellular oxygen conditions is critical to understanding host-pathogen interactions in the human gut.

Systems have been developed to address these gaps, each with a range of benefits and limitations ^21^. The widely available Transwell system has been adapted to maintain anaerobic bacterial viability by sealing the oxygenated basolateral media and placing the plate in anaerobic chamber ^22^. However, because the basolateral media is not oxygenated during the co-culture, the system can only be used for up to 8 hours. The Host-Microbiota Interaction (HMI) ^23^ and The Human Microbial Cross-talk (HuMiX) ^24^ models have also been developed and have been shown to support growth of anaerobic bacteria. HMI and HuMiX also have fluid flow, allowing co-culture for 24-48 hours. However, direct contact between bacteria and human cells is not possible. Additionally, a major limitation of these systems is the use of the Caco-2 cell line as opposed to a more physiologically-relevant cell culture model, such as human enteroids. Advancing on these systems, a recent study by Jalili-Firoozinezhad *et al*. (2018) demonstrated the ability to co-culture anaerobic bacteria with human enteroids and maintain viability for up to 120 hours ^25^. However, as with other microfluidics-based approaches, it is highly technical, labor-intensive, and proprietary.

Thus, there is a critical need for a cost-effective, easy to assemble system to model host-anaerobe interactions under physiologically relevant oxygen conditions. To address this gap, we developed a simple method for co-culturing obligate anaerobic bacteria with human intestinal enteroid monolayers under defined oxygen conditions.

## Materials and Methods

### Host-Anaerobe Co-Culture System, A Brief Description

Enteroids were seeded as monolayers on 6.5 mm transwell inserts (COSTAR 3470). These were placed into modified gaskets that were sealed in place using double-sided adhesive tape on a 24-well gas-permeable plate. The entire apparatus was kept in an anaerobic chamber (90% N_2_/5% H_2_/5% CO_2_) to allow growth of anaerobic bacteria on the apical surface. Gas (5% CO_2_/balance N_2_) containing oxygen (5.6% or 10.2%) was pumped from an external tank through the base of the plate to supply oxygen to the basolateral side of the monolayer. This constitutes a simple, cost-effective method for co-culturing obligate anaerobic bacteria with human, intestinal enteroid monolayers under variable oxygen conditions. A detailed description of system design, setup, and operation can be found in the supplementary materials provided (**Supplementary Material 1**).

### Human Jejunal Enteroids

Three different types of media were used to wash (complete medium without growth factors, CMGF-), proliferate (complete medium with growth factors, CMGF+), and differentiate/maintain (differentiation medium) human intestinal enteroids, initially produced from human intestinal biopsies and cultured as detailed by Saxena *et al*. (2016) ^8^. Three-dimensional cultures of jejunal enteroids were generated, cultured, and seeded as monolayers on 6.5 mm transwell inserts (COSTAR 3470). For monolayer formation, these dense 3D enteroid cultures were prepared as single cell suspensions using successive trypsin/EDTA incubation steps. Proliferating cultures were re-suspended in growth media (approximately 5×10^5^ cells / 100µL media) and seeded onto a diluted (1:40 in cold PBS) Matrigel/collagen coating film on the Transwell ^8^. This process allows enteroid monolayer formation. Growth medium was replaced with differentiation medium after 1-2 days, once trans-epithelial electrical resistance approached 300 Ω, indicating monolayer confluence. Monolayers were differentiated for 4 days prior to performing the co-culture experiments.

Specific enteroids used in each experiment are referred to by the intestinal segment of origin (J – Jejunum) followed by a number indicating donor origin (e.g., J2, J3, J8, J11 each originated from jejunum tissue from 4 different donors). Consent for the original donor tissue was obtained under IRB protocols H-13793 and H-31910 approved by the Baylor College of Medicine Institutional Review Board. Established lines were used in all experiments when between 10-15 passages in culture.

### Bacterial Strains, Preparation, and Quantification

Two commensal species, *Bacteroides thetaiotaomicron* (nanoanaerobe) and *Blautia* sp. (obligate anaerobe), cultivated from a healthy human microbiome were used in this study. These microbes were selected to represent the two most abundant phyla in the human gut ^26^. We utilized the continuous-flow mini-bioreactor array system described in Auchtung *et al.* (2015) to ensure that the bacterial cultures remained consistent between experiments ^27^. Continuous-flow culture models allow for studies to be performed during an extended time period under conditions where pH, nutrient availability, and washout of waste products and dead cells can be well controlled ^27^. Notably, the EACC system does not require mini-bioreactor arrays for operation and bacteria can be prepared according to user preference. Bioreactor medium was prepared as described in Auchtung *et al*. (2015) ^27^ and inoculated with our two bacterial isolates. After inoculation of the bioreactor, bacteria were allowed to equilibrate for 16–18 h prior to the initiation of flow at 1.875 ml/h (8-h retention time).

Prior to co-culture, approximately 0.2mL of bacterial culture was removed, serially diluted, and plated on bioreactor media agar plates (*B. thetaiotaomicron*) or GM-17 agar plates (*Blautia* sp.). These were cultured under anaerobic conditions for 24-48 hours to determine viability and concentration (CFUs/mL). Enteroid monolayers were inoculated with either *B. thetaiotaomicron* or *Blautia* sp. at 1×10^6^ CFU/mL or 1×10^5^ CFU/mL, respectively. *Blautia* sp. was inoculated at a lower concentration because of its considerably fast doubling time. At each experimentally defined timepoint, a 5µL sample was taken from the apical compartment of the Transwell, serially diluted, and spot plated in duplicate on the above described agar plates under anaerobic conditions to determine viability and concentration (CFUs/mL).

### Determination of Oxygen Concentration

Oxygen was measured with a 500 <u>u</u>M diameter Clark-type microelectrode (Unisense A/S) that was calibrated according to manufacturer’s recommendation under anoxic and atmospheric oxygen conditions. Measurements were taken at a consistent and defined position within the Transwell with the use of a micromanipulator to ensure precision.

### Simulation of Oxygen Transport in Anaerobic Model

A finite element simulation of oxygen transport in the EACC was constructed in COMSOL Multiphysics 5.3. Two-dimensional, radially symmetrical models of the gas-permeable, anaerobic Transwell system were drawn to scale, and a very fine tetrahedral mesh was used to construct a wireframe. Dimensions of the 24 well plate and Transwell insert were obtained from the manufacturer’s specifications. To simulate oxygen transport in the EACC with 5.6% O_2_, the concentration of O_2_ at the top and bottom of the well were defined to be 0.164% and 5.677%, respectively, as measured empirically by Clark electrode. Diffusion coefficients for oxygen were assumed to be 1.78 x 10^-6^ m^2^ s^-1^ in the membrane ^28^ and 3.0 x 10^-9^ m^2^ s^-1^ in the media ^29^. The semi-permeable Transwell membrane was modeled as a thin diffusion barrier with a diffusion coefficient equal to the diffusivity of the media multiplied by the porosity of the membrane (0.19).

Oxygen consumption by the enteroid monolayer was modeled using Michaelis-Menten kinetics:

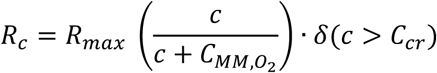

*R*_*c*_ is the rate of consumption, *R*_*max*_ is the maximum oxygen consumption rate, *c* is the concentration of oxygen, *C*_*MM,O_2_*_ is the Michaelis-Menten constant equal to the oxygen concentration at which oxygen consumption is one half of the maximal rate, and δ is a step-down function that stops oxygen consumption when its concentration drops below the threshold that can sustain survival in long-term cultures (*C*_*cr*_). Literature values were used for *C*_*MM,O_2_*_, 1.0 x 10^-3^ mol m^-3^ and *C*_*cr*_, 1.0 x 10^-4^ mol m^-330^.

*R*_*max*_ was determined experimentally by culturing enteroid monolayers on collagen type I gels (Rat Tail Collagen Type I, Corning) in an oxygen-sensing well plate (OP96C, PreSens Precision Sensing GmbH) for 5 days in a standard tissue culture incubator. In this setting, *R*_*c*_ approximates *R*_*max*_ due to the high O_2_ concentration. The equilibrium O_2_ concentration of the monolayers was measured, and *R*_*max*_was calculated using a finite element model of the well plate.

### Trans-epithelial Resistance

The barrier integrity of enteroid monolayers was determined with trans-epithelial electrical resistance (TEER) as measured with an epithelial Volt/Ohm meter (Millipore MERS 000-01).

### Trypan-Blue Dye Exclusion Assay for Cell Survival

To quantify cell survival, enteroid monolayers were treated with 200µL 0.05% Trypsin/EDTA, incubated at 37°C for 5 minutes, and suspended into a single-cell homogenate. The resulting suspension was mixed with an equal volume of Trypan Blue, loaded into a dual chamber counting slide (BioRad 145-0011) and quantified using a TC20 Automated Cell Counter (BioRad 1450103).

### Histology, immunofluorescence staining, and imaging

To preserve the mucus layer and maintain bacterial attachment, enteroid monolayers were fixed in Carnoy’s fixative (Election Microscopy Science 64130-50) at room temperature for 4 hours, embedded in paraffin, sectioned, and stained with Hemotoxylin and Eosin. Slides were also stained with Alcian Blue to identify the mucus layer and discriminate goblet cells ^8^.

### Bacterial Fluorescence In Situ Hybridization

Tissue was fixed in Carnoy’s fixative at room temperature for 4 hours, then embedded in paraffin wax. 4um sections were mounted on glass slides, baked at 60°C for 1 hour, then de-paraffinized with xylene and dehydrated in series from 50% to 100% ethanol. The probe used was a previously validated 5’ Alexafluor488-labelled universal bacterial probe (Uni519; 5’-GTATTACCGCGGCTGCTG-3’) targeted to 16S rRNA ^31^. Samples were counterstained with DAPI. The probe was hybridized to the samples by adding 15µL of the 2µM probe to each slide and placing in a 45°C hybridization chamber for 45 minutes. Slides were imaged using a Nikon Eclipse 3000 microscope.

### Gene Expression Profiling

The Qiagen RNeasy Mini-Kit was used to extract RNA from enteroid monolayers. This was followed by gDNA elimination and cDNA conversion via the Qiagen RT2 First Strand Kit (330401). Gene expression was evaluated using the RT2 Profiler PCR Arrays for human tight junctions (330231) and human antibacterial response (330231) using the associated RT2 SYBR Green ROX qPCR Mastermix (330523). Twelve samples (3 patient lineages, 4 experimental conditions), each from 2-3 pooled enteroid monolayers, were run in duplicate on the ABI ViiA 7. Cycle Cut-off was established at a CT of 37 for inclusion in analysis. All expression data was normalized to the 5 housekeeping genes included on the array and all treatment groups were normalized to the cell-line specific incubator control on each RT2 plate. Each RT2 plate ran all four treatment groups within a single enteroid lineage to eliminate variability between runs. All samples passed all quality control metrics for PCR array reproducibility, RT efficiency, and genomic DNA contamination. All gene lists were stripped of type C errors (if 1/12 samples fell below CT cut-off), and genes with >2 difference in fold expression between replicates of any sample were excluded. Of the 169 genes analyzed, 107 were retained in the final analysis. A 1.5 difference in fold change in expression was the minimum threshold for “differential” regulation.

## Results

### Assembly and validation of the EACC

After 4 days of differentiation, seeded Transwells were fitted into the modified gaskets of the EACC and sealed in place using double-sided adhesive tape (D969PK Silicone/Acrylic Differential Tape by Specialty Tapes Manufacturing), on a 24-well plate with a gas-permeable base (**Figure 1a, 1b**). Blood gas (5% CO_2_ / 5.6% or 10.2% O_2_ / Balance N_2_) was pumped from an external tank through the base of the plate to feed the basolateral side of the monolayer, which was placed in differentiation media. A mixture of differentiation media and bioreactor media (300μL of a 1:1 mixture) was added to the apical compartment (**Supplementary Figure 1a**). Dissolved oxygen concentration, as measured by a clark-type microelectrode, reflected a steep oxygen gradient across the monolayer (**Figure 1c)**. Following 24 hours of culture, measured dissolved oxygen in the apical compartment was effectively zero while the oxygenation of the basolateral compartment reflected delivered blood gas oxygen concentration (5.6% or 10.2% oxygen). In a cellular model of oxygen gradient across the epithelium, distribution of oxygen within the system in the presence of an enteroid monolayer demonstrates that enteroids scavenge residual apical oxygen and prevent oxygen leaking into the apical compartment (**Supplementary Figure 1b**). Using this model and considering the oxygen consumption of an enteroid monolayer, the calculated time to equilibrium at all apical points above the Transwell was 2 hours (**Supplementary Figure 1c**). This model predicts the time at which the entire system equilibrates and anaerobic bacteria can be added to the apical compartment.

**Figure 1.**
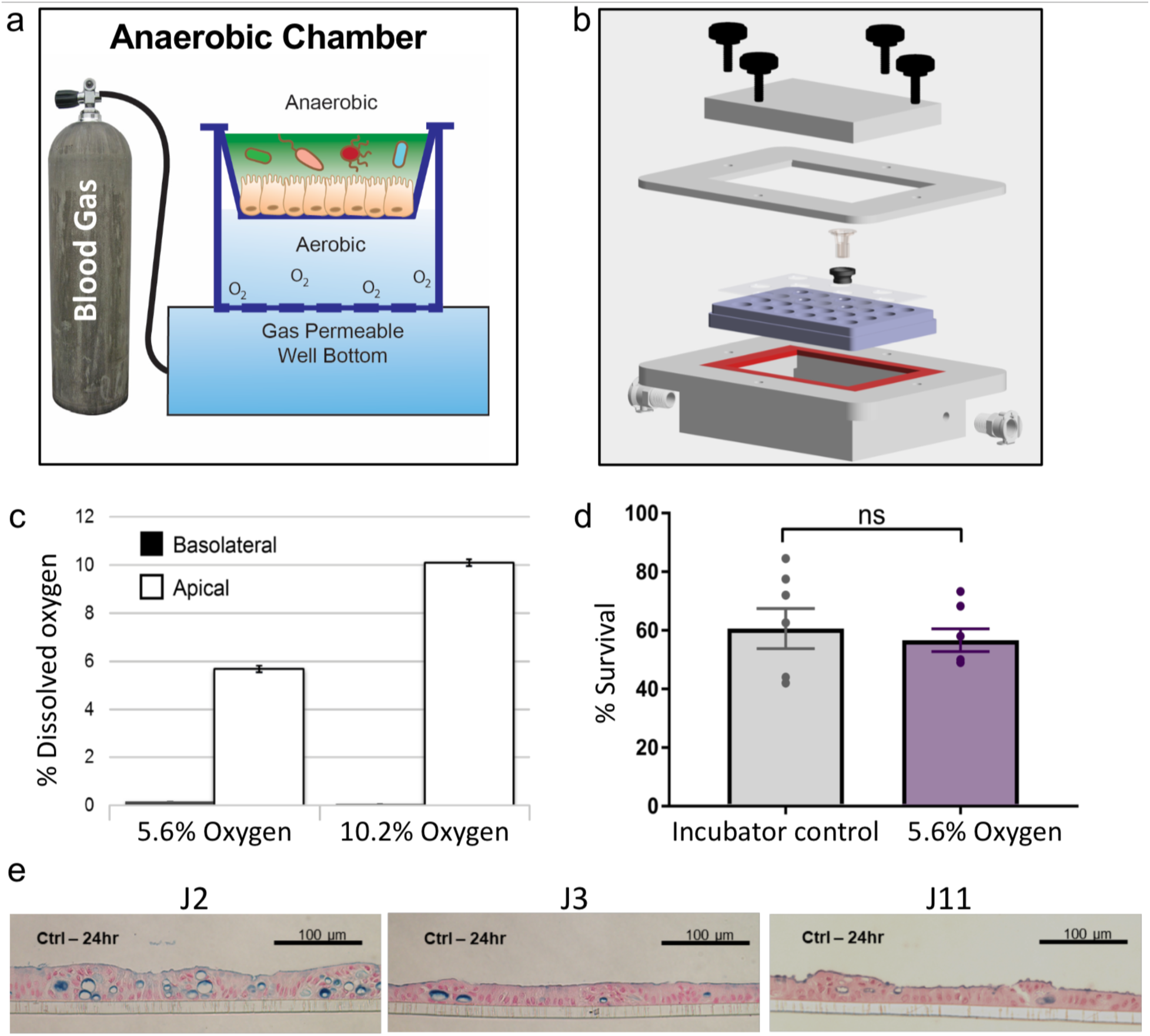
Development and validation of the Enteroid-Anaerobe Co-Culture (EACC) system reflects in vivo oxygen gradients, morphology and survival. (a) In the conceptual system assembly, the entire assembly is maintained within a standard anaerobic chamber. (b) Within the assembly itself, Transwells seeded with enteroid monolayers were fitted into modified gaskets and sealed in place, using a special double-sided adhesive tape on a 24-well plate with a gas-permeable base. Blood gas was pumped from an external tank through the base of the plate to feed the basolateral side of the monolayer, while anaerobic bacterial are grown in the anaerobic apical compartments. (c) Experimentally measured dissolved oxygen concentration demonstrates consumption of oxygen by enteroid monolayer resulting in an anaerobic apical environment when exposed, basolaterally, to blood gas with either 5.6% or 10.2% oxygen. (d) Enteroid monolayers survive exposure to basolateral physiological hypoxia as measured by Trypan Blue dye exclusion (N=7/group). (e) Enteroid monolayers are also polarized and have normal morphology as measured by Alcian Blue staining.

There was no significant difference in enteroid survival between enteroid monolayers cultured in standard incubator conditions (incubator controls; ICs) and those cultured under physiologic hypoxia (5.6% oxygen) within the EACC system over 24 hours (**Figure 1d**). Alcian Blue stain of monolayers at 5.6% oxygen after 24 hours showed normal cellular morphology and the formation of a mucus layer with appropriate polarity (**Figure 1e**).

### Expression of hypoxia-associated phenotypes

TEER, a proxy for epithelial barrier integrity, increased in response to 24 hours of 5.6% basolateral oxygen exposure within the EACC system relative to ICs (**Figure 2a**). In the literature, 5.6% oxygen is reported as more reflective of physioxia than inflammatory hypoxia as it is sufficient to promote physioxia-associated phenotypes without resulting in substantial HIF1a expression ^20^. This finding was observed in each patient derived line (J2, J3, J8, and J11), as well as in the traditionally used CaCO2 cell line (**Supplementary Figure 2a**). Studies have shown that low oxygen conditions are critical for the constitutive expression of innate immune factors and the expression of genes that enable epithelial cells to function as an effective barrier ^32–34^. Concordantly, we screened three of the jejunal lines (J2, J3, and J11) for expression of 168 genes associated with epithelial barrier integrity, innate immunity, and anti-microbial response. Of the 107 genes that passed our filtration criteria, 32 were significantly upregulated in 5.6% oxygen relative to ICs and 3 genes were significantly down-regulated (**Figure 2b**). Specifically, 21 barrier integrity genes are significantly upregulated (**Figure 2c**) across all three patient lines, corroborating the TEER phenotype (**Figure 2a)**. Physiological hypoxia further upregulated 12 anti-microbial response genes, including TLR4, TLR6, and MYD88 (**Figure 2e**). As expected, the oxygen-sensitive transcription factor hypoxia-inducible factor-1alpha (HIF-a) was also increased. IL1 (CXCL1), IL8 (CXCL8) and TLR2 were all significantly down-regulated in response to hypoxia (**Figure 2d**). Heatmap analysis illustrated the significant and somewhat patient-specific changes in gene expression of both epithelial barrier integrity genes and anti-microbial immune response genes during physiological hypoxia (i.e., 5.6% oxygen) (**Supplementary Figure 3)**.

**Figure 2.**
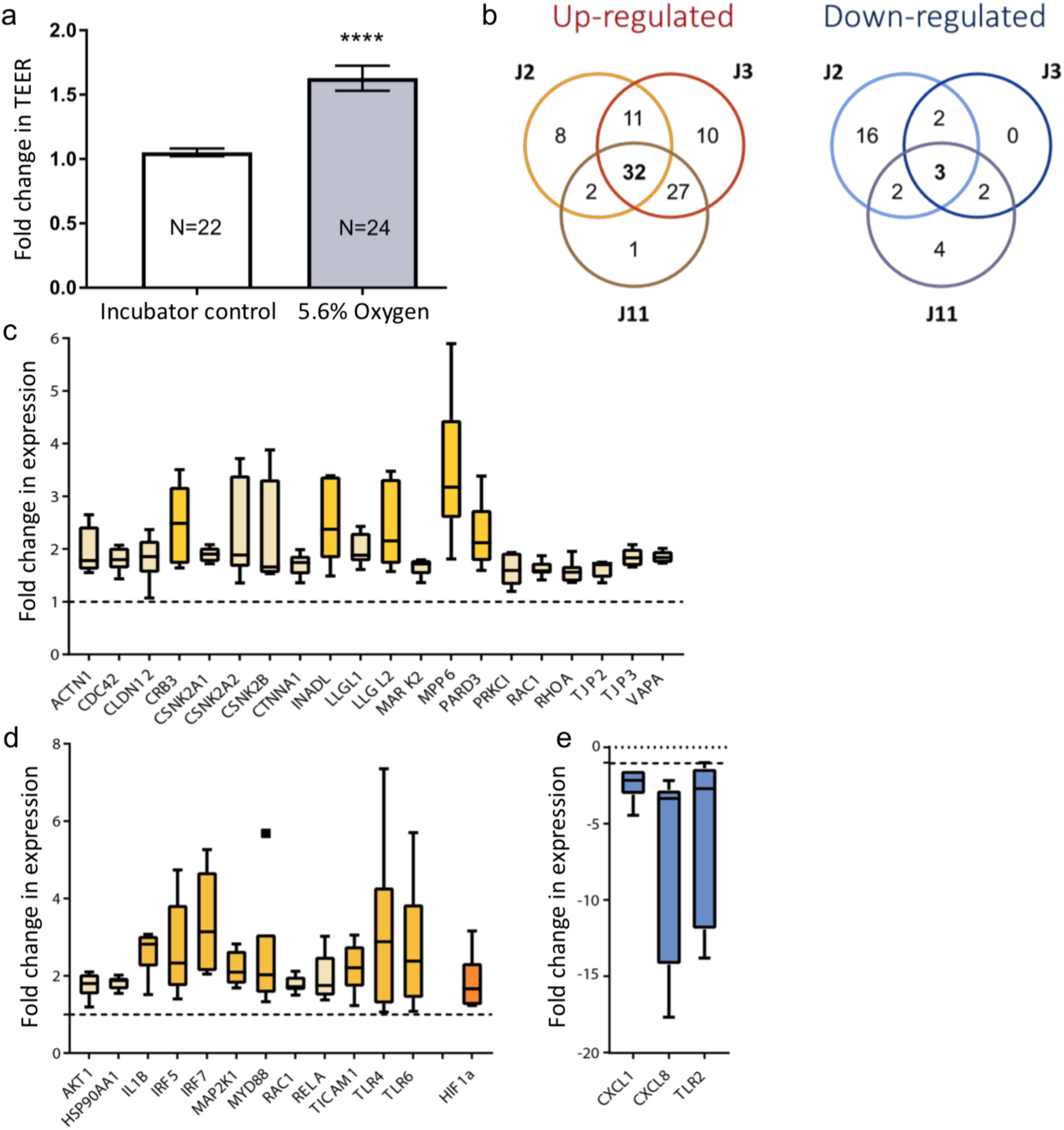
Exposure to 5.6% basolateral oxygen within the Enteroid-Anaerobe Co-Culture (EACC) system reflects expression of physioxia-associated phenotypes. (a) Increased trans-epithelial electrical resistance (TEER) in 5.6% basolateral oxygen. (b) Venn diagrams showing consistent gene signature related to hypoxia across 3 jejunal lines, 32 genes were upregulated relative to standard incubator condition and 3 were down-regulated. (c) Barrier integrity genes that were consistently upregulated in all 3 jejunal lines in response to hypoxia, which included Junction Proteins (INADL, PARD3, TJP2/3) and Cell Surface Receptors (CRB3, CLDN12). (d) Core anti-microbial response genes consistently up-regulated in 3 jejunal lines in response to hypoxia. (e) Core anti-microbial response genes consistently down-regulated in 3 jejunal lines in response to hypoxia.

**Figure 3.**
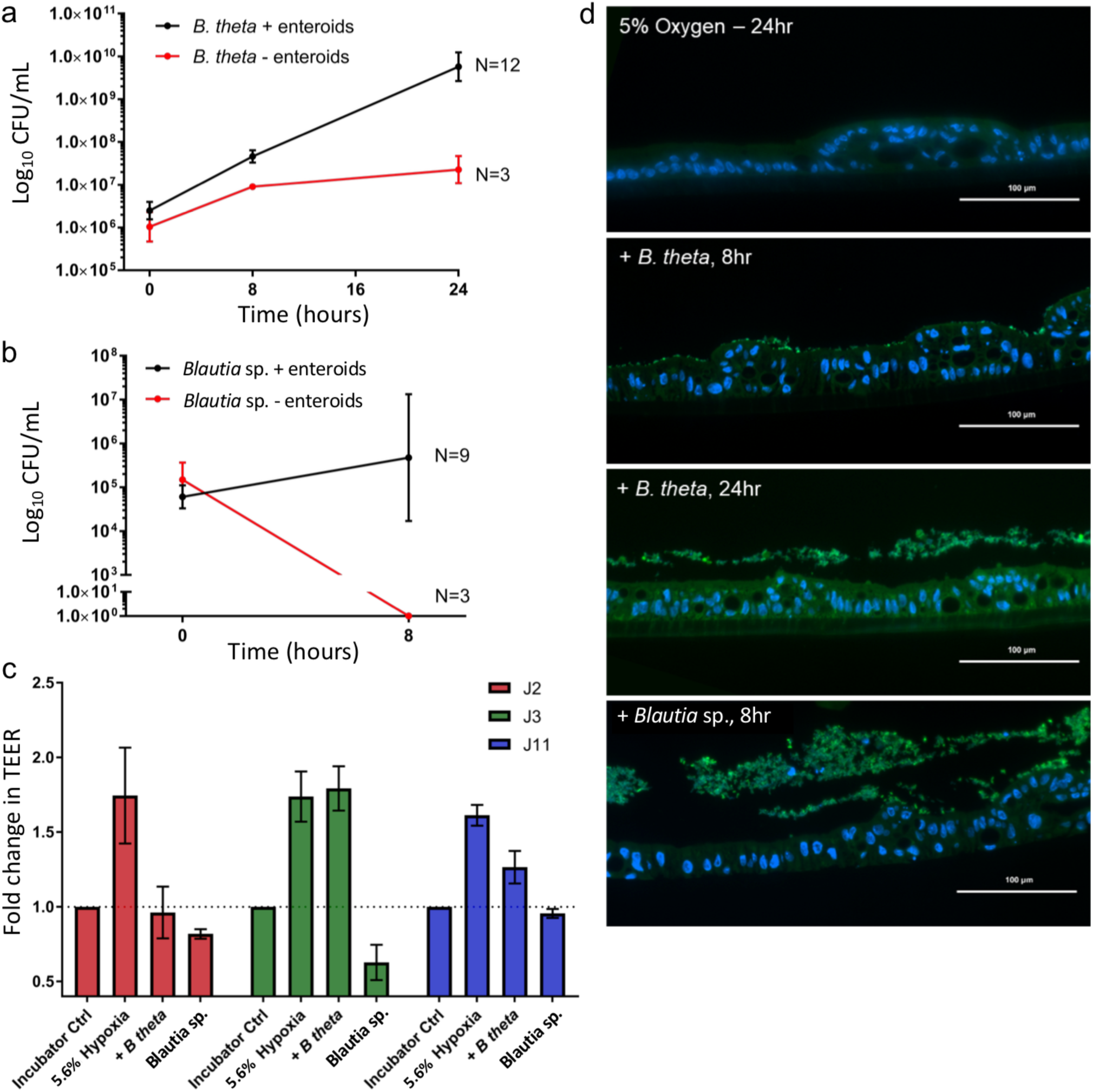
The Enteroid-Anaerobe Co-Culture (EACC) system supports co-culture of enteroid monolayers with anaerobic bacteria for up to 24 hours. (a) The EACC system supports growth of *B. thetaiotaomicron (B. theta*) for at least 24 hours. (b) The Co-Culture system also supports the growth of obligately anaerobic *Blautia sp. f*or at least 8 hours. Because *Blautia* cannot survive oxygen exposure, CFU/mL was zero in the absence of enteroid monolayers. (c) The effect of bacterial co-culture on enteroid trans-epithelial resistance is dependent on enteroid lineage (n=7 for Ctrl, Hypoxia, and *B. theta* treated groups; n=3 for *Blautia* treated groups). (d) Fluorescent In-Situ Hybridization for the 16S rRNA gene reflects bacterial presence and growth in green.

Gene Ontology analysis highlighted several significantly activated pathways, including nitric oxide biosynthesis, NFkB activation, PRR signaling, IL6 production, positive regulation of cell communication, inflammatory response activation, and maintenance of cell polarity (**Supplementary Table 1**). Under higher oxygen conditions, the cellular HIF-1α level is regulated by hydroxylation by prolyl hydroxylases (PHDs) which regulate HIF-1α. During hypoxia, endogenous nitric oxide disables PHDs, allowing HIF-1α to accumulate and activate its downstream gene targets ^14^. Thus, positive regulation of nitric oxide biosynthesis as the top activated pathway acts as a secondary validation that the response observed is, in fact, a response to physiological hypoxia.

### Enteroid-Anaerobe Co-Culture

*Bacteroides* (Bacteroidetes phylum) and Lachnospiraceae (Firmicutes phylum) are among the most abundant bacteria in the human gut ^26,35^. *Bacteroides thetaiotaomicron* is a gram-negative, acetate-producing nanoanaerobe from the Bacteroidetes phylum ^36^. Member of the *Blautia* genus (Lachnospiraceae family) are gram-positive, lactate- and acetate-producing obligate anaerobes from the Firmicutes phylum and have been associated with reduced death from graft-vs-host disease ^37^. To test the EACC system, we co-cultured *B. thetaiotaomicron* and *Blautia* sp. with human jejunal enteroid monolayers under 5.6% basolateral oxygen. Here, our experimental approach was similar to that of earlier hypoxic experiments. However, in co-culture experiments, following 2 hours of equilibration within the anaerobic chamber, approximately 3×10^4^ bacteria were inoculated into the apical compartment (**Supplementary Figure 4**). Bacteria were co-cultured in the presence and in the absence of enteroid monolayers to validate the model that enteroids consume residual apical oxygen, making the upper compartment effectively anaerobic. The EACC system supported the growth of the nanoanaerobe *B. thetaiotaomicron*, which had a slow doubling time and is able to survive in low oxygen conditions, for at least 24 hours as measured by colony forming units (CFU/mL) (**Figure 3a**). *B. thetaiotaomicron* co-culture reduced TEER in an enteroid-specific manner (**Figure 3c**).

**Figure 4.**
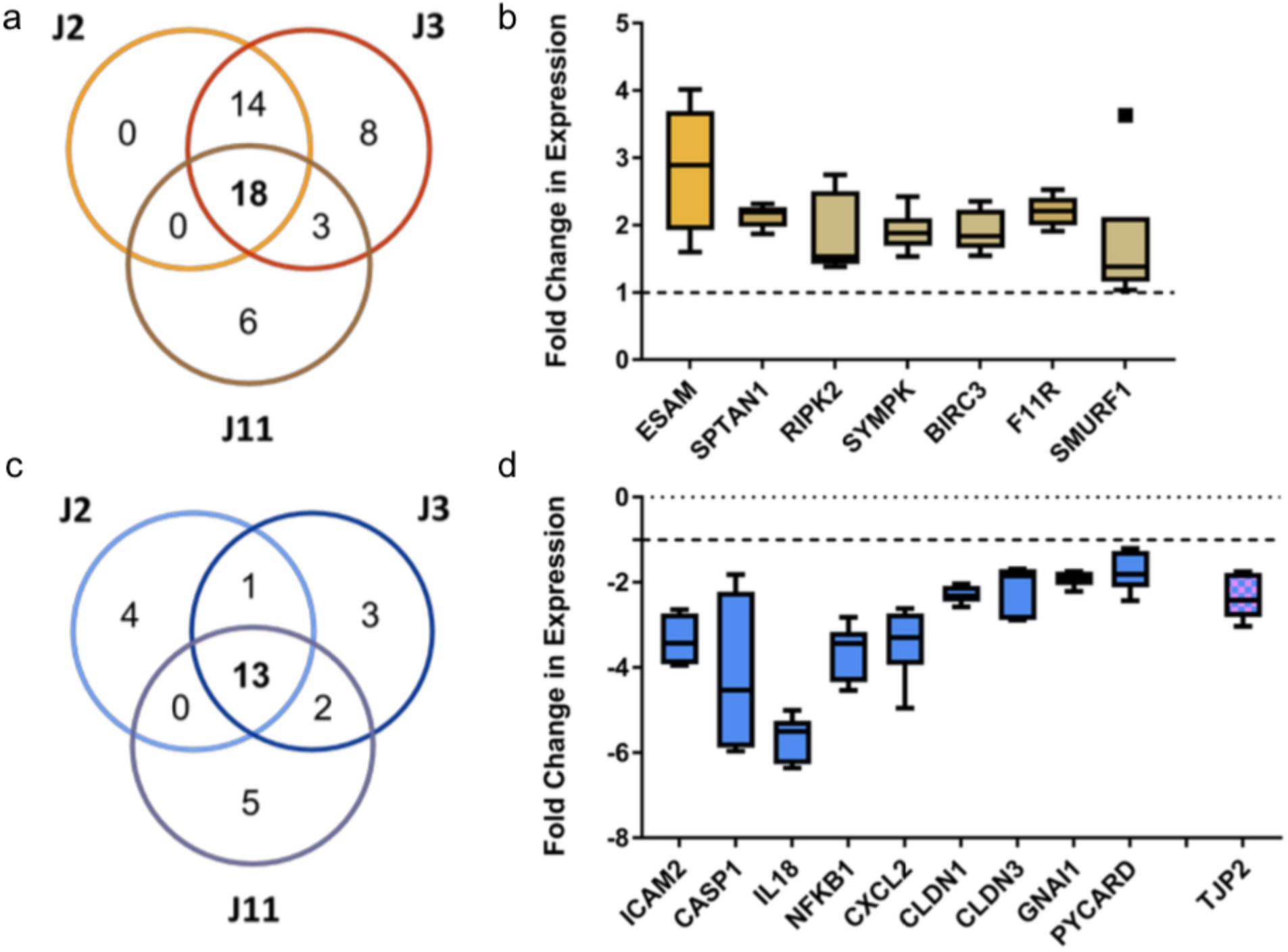
Effects of Bacterial co-culture on gene expression. (a) 18 genes were consistently upregulated across all three jejunal lines during *B. thetaiotaomicron (B. theta*) co-culture relative to standard incubator condition, 11 of which were also upregulated during hypoxia. (b) Of the genes consistently and specifically upregulated include ECAM, which is involved in trans-endothelial leukocyte migration. (c) 13 genes were consistently down-regulated during *B. theta* co-culture hypoxia compared to standard incubator conditions, many of which associated with immune response (IL18, NFkB, CXCL2, etc) and intestinal permeability (CLDN1/3, TJP2).

The EACC system also supported the survival and growth of *Blautia* sp., which is not able to tolerate any residual oxygen (**Figure 3b**). The survival and growth of *Blautia* was dependent on enteroid lineage, as observed by the large error bars (**Figure 3b**). As expected, in the absence of an enteroid monolayer the basolateral oxygen diffused into the apical media and was lethal to the *Blautia* sp. (**Figure 3b**). Similar to *B. thetaiotaomicron* co-culture, *Blautia* affected TEER in a patient-specific manner (**Figure 3c**). Fluorescent in-situ hybridization for the 16S rRNA gene supported bacterial presence and growth (**Figure 3d**). Interestingly, *B. thetaiotaomicron* adheres to the monolayer at 8 hours and is subsequently detached from the epithelium by 24 hours, suggesting epithelial remodeling and increased mucus production.

### Transcriptomic Response to Anaerobic Co-Culture

Controlled by the oxygen-sensitive transcriptional regulators HIF-1α and NFkB, hypoxia/physioxia alter the intestinal epithelium in a variety of ways that impact bacterial invasion and host-microbe interactions ^15,38^. Hypoxic tissue also features enhanced expression of barrier protective genes and mucins that limit bacterial translocation, release antimicrobial factors, and recruit inflammatory cells ^13^. Selective remodeling of cell membranes further hinders bacterial attachment ^39^. To determine whether enteroids within the EACC system recapitulate the features of a physiologically hypoxic epithelium, we assessed the transcriptomic response of human enteroids to anaerobic co-culture with *B. thetaiotaomicron* or *Blautia* sp. in the presence of 5.6% basolateral oxygen.

Heatmap analysis illustrated the significant and highly patient-specific changes in gene expression of both epithelial barrier integrity genes and anti-microbial immune response genes following bacterial co-culture (**Supplementary Figure 5**). All three enteroid lines showed a slight reduction of the hypoxic phenotype when co-cultured with *B. thetaiotaomicron*. However, gene expression in response to *Blautia* co-culture varied between enteroid lines. In response to *B. thetaiotaomicron* co-culture, 18 genes were significantly upregulated across all three patient lines relative to ICs, 11 of which were also upregulated during physiological hypoxia indicating a consistent carry-over of the hypoxic phenotype (**Figure 4a**). However, 7 additional genes were shown to be upregulated in response to co-culture independent of response to hypoxia (**Figure 4b**). Most of the upregulated genes were responsible for regulating barrier integrity, including cytoskeleton regulation (SPTAN1, SMURF1) and junction maintenance (ESAM, SYMPK, F11R). Substantially more genes (13) were down-regulated in response to *B. thetaiotaomicron* co-culture as compared to ICs (at ambient oxygen), and the three genes that were downregulated during physiological hypoxia (IL1, IL8, and TLR2) were also downregulated during hypoxic co-culture **(Figure 4c)**. Interestingly, genes responsible for inflammatory response (ICAM2, IL18, NFKB1, IL2) and induction of apoptosis (CASP1, PYCARD) are strongly downregulated during co-culture, suggesting an immunomodulatory role for *B. thetaiotaomicron* in the gut **(Figure 4d)**. Many of the genes significantly upregulated during hypoxia were mitigated to IC levels of expression following *B. thetaiotaomicron* co-culture (**Supplementary Figure 6**).

Gene Ontology analysis highlighted several significantly activated pathways, including regulation of cell proliferation, inflammatory response, interspecies interaction between organisms, activation of immune response, cytokine production, NFkB activation, and response to a molecule of bacterial origin (**Supplementary Table 2**). These findings indicate that much of the original hypoxic gene expression phenotype remained during co-culture. However, there was little overlap in gene expression profiles between jejunal lines to the same treatment group, suggesting patient-specific responses to bacteria.

## Discussion

Physiologically normal oxygen concentrations in the intestinal mucosa range from 1-11% oxygen. Yet most in-vitro experimentation is performed in incubator conditions (~18.8% oxygen), making these cultures effectively hyperoxic and their results potentially misleading ^13,19^. Our findings are consistent with resolution of a hyperoxic phenotype into physioxia, in that we observe increased barrier integrity at 5.6% basolateral oxygen, increased expression of barrier and antimicrobial response genes, and decreased production of the proinflammatory cytokines IL-8 and IL-1. Interestingly, inflammatory hypoxia is also known to upregulate IL-8 production, which suggests that the EACC system at 5.6% oxygen more closely resembles physioxia than inflammatory hypoxia ^16,40^. From pathway analysis, we also observed positive regulation of NFkB and nitric oxide biosynthesis. Nitric oxide, a gasotransmitter, controls HIF-1α stability during physioxia and inflammatory hypoxia ^14^. Within the EACC system, induction of physiological hypoxia has crucial implications for downstream co-culture studies. For example, increased expression of TLR4 coupled with decreased expression of TLR2 suggests increased sensitivity to gram-negative bacteria and decreased sensitivity to gram-positive bacteria, respectively. We also observed upregulation of several genes associated with the type 1 interferon response (IRF5, IRF7, MyD88, etc.) which regulates cellular response to viral infection. Furthermore, increased barrier integrity discourages bacterial translocation into the basolateral space. Epithelial response to physiologically normal hypoxic conditions alters baseline cellular response to both bacteria and viruses, demonstrating the need to address physiological and inflammatory hypoxia in host-microbe interaction studies. Furthermore, these findings suggest that co-culture of epithelial cells with other cell types, such as immune cells, may yield confounding results via the introduction of oxygen consuming cells and the subsequent induction of cellular hypoxia.

The method of tissue culture can influence partial oxygen pressure. For instance, when seeded as monolayers on the base of a standard 24-well cell culture plate, the ppO2 of CaCO2 cultures can fall as low as 30 mmHg (~4% oxygen). However, when grown on membrane inserts or at the base of standard gas-permeable plates, ppO2 generally bottoms out at 125mmHg (~16.5% oxygen) ^18^. Hyperoxia is known to induce IL-8 production in some cell types as an overabundance of luminal oxygen is frequently indicative of opportunistic or pathogenic bacterial invasion ^41,42^. During hyperoxia, barrier integrity is diminished to permit luminal antigen sampling by dendritic cells and to allow immune cells to infiltrate the lumen. Intestinal epithelial cells produce IL-8 in order to recruit polymorphonuclear leukocytes (PMNs), such as neutrophils, to the site of infection ^16^. Once present, PMNs undergo an oxidative burst that consumes 10x more oxygen than any other cell in the body, depleting the epithelial environment of residual oxygen and promoting serosal hypoxia. Reduced serosal oxygen then activates HIF-1α and canonical NFkB, which are both oxygen sensitive master regulators whose downstream targets increase barrier integrity, boost anti-microbial peptide production, and prompt membrane/cytoskeletal reorganization.

The intestinal epithelium serves as the first line of defense against invasive and opportunistic pathogens and is critical in the establishment of tolerogenic response. Both commensal and pathogenic species have been shown to regulate oxygen pressure in the mucosa and influence hypoxia-inducible factors signaling in enterocytes ^43^. In addition to being responsive to oxygen, NFkB and HIF-1α are also regulated by inflammatory mediators, such as cytokines, and by bacterial products, such as Lipopolysaccharide (LPS). LPS makes up the outer membrane of gram-negative bacteria, including that of *B. thetaiotaomicron*. However, while *Bacteroides* produces LPS, it does not elicit a potent pro-inflammatory response because its protein structure differs significantly from pathogen-derived LPS ^36^. *B. thetaiotaomicron* is generally recognized as a commensal in the gut and is known to stimulate secretory lineage differentiation, mucin glycosylation, and decrease enteroendocrine cell activity ^44^. Our findings reflect those reported in the literature, as co-culture with *B. thetaiotaomicron* generally ablates the upregulation of many anti-microbial (e.g., IL18, Caspase 1, NFkB1) and tight junction (e.g., CLDN1, CLDN2, TJP2) response genes and demonstrates down-regulation of inflammatory genes (e.g., IL-8, IL-1, MyD88, and TLR-2). Fluorescent In-Situ Hybridization staining for the 16S rRNA gene demonstrates the attachment of *B. thetaiotaomicron* at 8 hours and repulsion of the bacteria at 24 hours, suggesting a reorganization of the cell surface to hinder bacterial attachment, likely through increased production of mucus and antimicrobial peptides. However, the findings associated with *Blautia* co-culture suggest that cellular response to this beneficial bacterium may be patient-dependent. An abundance of *Blautia* is associated with protection from graft-vs-host disease and is currently in clinical trials for irritable bowel syndrome ^37^. Studies have indicated that host genetics, as well as environmental exposure history, can influence epithelial response to both pathogens and commensals. Thus, it is possible that a microbial therapeutic such as *Blautia* may also have patient-specific efficacy, demonstrating the need for a personalized medicine approach in host-microbe interaction studies.

The EACC system described herein permits the investigation of host-microbe interactions with precision control of basolateral oxygen concentration. Given our oxygen consumption models, the operating range of this system is approximately 2%-10% oxygen. These values more closely model oxygen concentrations found *in vivo*, which can range from 1% in inflamed tissues and tumors to 11% during meal digestion. This system reaches steady-state equilibrium within two hours of setup and accurately models the steep oxygen gradient found in-vivo. Studies of hypoxia in epithelial cell lines show that HIF1a levels generally peak between 2 and 6 hours. We also show that, at 5.6% basolateral oxygen, we induce many of the hallmarks of hypoxia including HIF-1α expression, increased barrier integrity, upregulation of tight-junction and anti-microbial response genes, activation of the NFkB and NO biosynthesis pathways, and reduced production of pro-inflammatory cytokines such as IL-8. Thus, this system is optimal for elucidation patient-specific responses to serosal oxygenation.

Importantly, the EACC system facilitates short term co-culture of commensal nano-anaerobes (*B. thetaiotaomicron*) and obligate anaerobes (*Blautia* sp.) with enteroid monolayers. Both enteroid monolayers and anaerobes survived co-culture and responded reproducibly in a predicted manner. Other host-microbe co-culture systems exist within the field, as summarized by Martels *et al*. (2017), and each provides a range of benefits and challenges ^45^. Microfluidics-based systems offer flow, known to stimulate differentiation and physiologically relevant phenotypes, but are generally IP-protected, highly technical, expensive, and difficult to scale-up. Furthermore, microfluidic devices are generally constructed from PDMS, an oxygen-porous material that threatens the survival of true obligate anaerobes. Notably, a recent study using microfluidic chips reports the ability to co-culture Caco-2 and primary cell lines with anaerobic bacteria, specially *Bacteroides fragilis* ^25^. While this is a major advancement in microfluidic systems, the dependency for using a specifically engineered chip for study limits throughput, increases the expense, and restricts the use of the system to specialist labs. Conversely, the EACC system presented here is constructed from common laboratory and commercial materials that are readily available at relatively low costs. Because it is based on the Transwell system, it is easily integrated into existing cell-culture laboratories and provides an affordable, high-throughput system for interrogating host-microbe interactions. The user is also able to use existing infrastructure, such as existing anaerobic chambers, with minor modification in some instances to allow for tubing carrying oxygen to the console to run in/out of the chamber. Furthermore, its simple assembly permits direct interaction between human and microbial cells and allows for temporal sampling or replenishment of the apical media if desired by the user. The EACC is further compatible with well-validated cell culture lines (e.g., Caco-2) and can be used for co-culture or specific isolates or a consortium of isolates, as well as entire sample types (e.g., stool samples). While outside the scope of the current study, the consistent oxygen delivery should allow for the introduction of immune cells into the system.

In spite of its strengths, the EACC as used in this study also has several limitations. First, the system is static, meaning that enteroid viability may be compromised by bacterial overgrowth and/or changes in media pH. However, this limitation could likely be overcome through use of slower growing bacteria or when the user replenishes media and/or dilutes bacterial cells to prevent overgrowth. Furthermore, we were able to observe significant changes in the host response to microbes within our 8-24 hour time frames, and the shorter experimentation time allowed more efficient experimental throughput. Second, owing to the necessity to create an airtight seal between the basolateral and apical compartments, it is not possible to continually monitor the TEER. However, we believe that ability to being microbial and host cells into direct contact, while maintaining the viability of both species, outweighs this disadvantage, especially when TEER measurements can be obtained at the start and end of the experiment in the EACC system. Finally, an inherent limitation of enteroids and related systems derived from cell lines is that lack of circulating immune cells in the model, although methods exists to include these in Transwell monolayers ^46^. Future work should aim to incorporate these into models, with the EACC system providing a platform that should allow this integration of microbial-epithelial-immune cells into a single robust system.

## Conclusion

The Enteroid-Anaerobe Co-Culture (EACC) system is constructed from commonly available, affordable materials and requires minimal technical expertise for assembly and operation. It recapitulates the steep oxygen gradient seen in vivo, induces expression of hypoxia-associated phenotypes, and sustains host-anaerobe interactions for at least 24 hours. We also demonstrate that enteroid monolayers exposed to physioxia, as compared to incubator levels of oxygen, mount significant barrier-integrity and anti-microbial responses, even in the absence of bacterial co-culture. It is paramount that basic and therapeutic investigation of host-microbe interactions be conducted under oxygenation conditions that are physiologically appropriate to the intestinal location and disease state. Furthermore, we show substantial patient-specific differences in gene expression responses to co-culturing enteroids with anaerobic bacteria, reinforcing that this system can serve as a personalized medicine approach to biotherapeutic screening and efficacy investigation.

## Declarations

### Consent for publication

N/A

### Availability of data and materials

The datasets used and/or analyzed during the current study are available from the corresponding author on reasonable request.

### Author Contributions

T.F, J.M.A, and J.F.P contributed to conceptualization of the research; T.F and J.M.A designed the research methodology; T.F, C.J.S, J.M.A, and R.L.W, performed the research; R.B, G-A.J, M.K.E, and J.F.P provided resources; T.F and R.L.W analyzed the data; T.F, C.J.S, and R.L.W wrote the manuscript; T.F, C.J.S, J.M.A, R.L.W, M.K.E, and J.F.P. reviewed and edited the manuscript. All authors read and approved the final manuscript.

## Acknowledgements

not applicable

