## Supplementary for "A novel human enteroid-anaerobe co-culture system to study microbial-host interaction under physiological hypoxia"

**Supplementary Material 1.** Detailed protocol for Enteroid-Anaerobe Co-Culture (EACC) system assembly.

### Preparation (Day before)

- Autoclave the gaskets, lid (if necessary), and test tubes lids (or other holder) per console

***Carry out all steps below in a sterile tissue culture hood***

### Setting up the console

- Place the magnetic stirrer in the base of the console
- Open the gas permeable tissue culture plate.
  - Take care not to touch or indent the gas permeable membrane on the base of the plate
- Using a 10ml syringe filled with vacuum grease and with 200ul pipette tip attached to aid accurate placement, apply vacuum grease to the bottom skirting of the gas permeable tissue culture plate to make ensure an air-tight seal and prevent leaks
- Plate the gas permeable tissue culture plate base onto the central rubber seal (red)
- Remove the lid from the gas permeable tissue culture plate
- Place the lid of the console over the top of the gas permeable tissue culture plate
- Tighten the console by hand using the screws
- Stick the plastic binding surface of the Silicone-Acrylic Differential Tape (SAD tape) on top of the gas permeable tissue culture plate
  - Cover half the plate then repeat for the other side. Make sure the tape doesn't overlap otherwise it wont be possible to remove the plastic to reveal the sticky surface
- Using a blade trim off the excess SAD tape from each side of the gas permeable tissue culture plate
- Taking care to not pierce the lower gas permeable layer of the gas permeable tissue culture plate (don't put blade too far into the well) and use a scalpel carefully carve out the desired wells (every other well) of the gas permeable tissue culture plate
  - Using the edge of the well to guide the blade in a circle
  - Use a checked setup (open every other well) to ensure the gaskets fit
- Pipette 600ul of differentiation media into each open well
- Return the lid of the gas permeable tissue culture plate

### Preparation of the gasket/Transwell

- Remove film from top layer of the SAD tape to expose the rubber binding surface of the adherent tape
  - Use common electrical tape to wrap around the outside of the plate – this will create a better seal for the lid and prevent the apical media evaporating.
- Set up the test tubes lids (or other suitable mount) in line to act as mounts for applying the gasket to the Transwell
- Using sterile forceps transfer a gasket to each test tube (lid).
- By hand, remove Transwell from the 24-well plate and slowly push the Transwell into the gasket
  - Keep 24-well plate for later when doing the TERs
- By hand, add each gasket/Transwell in turn to the gas permeable tissue culture plate

    - Press firmly down to ensure strong binding between red adhesive tape and rubber gasket
- Add custom lid (see separate methods) to the top of the gas permeable tissue culture plate
    - Standard gas permeable tissue culture plate can be used but need to be stuck down

***Carry out all steps below in the anaerobic hood***

- Attach the gas tube to the inlet and outlet ports
- Place console on magnetic stir plate and turn onto a low/medium spin
- Open the cylinder to turn on the flow from the blood gas into the console
- Set the gas regulator to the left to flow rate 15 for 30 seconds to purge the chamber
- Then turn the knob to the right to set the long-term flow rate to 0.5 for the remainder of the experiment.
    - Allow system to equilibrate for 2 hours before adding bacterial to the apical media

- Remove apical media
- Add 200ul 1:1 BRM and differentiation media to the apical side of the monolayer

### SUPPLEMENTARY TABLES

**Supplementary Table 1. The top gene ontology pathways induced in response to physiological hypoxia.**

**Supplementary Table 2. The top gene ontology pathways induced in response to *B. theta* co-culture under physiological hypoxia**

### SUPPLEMENTARY FIGURES

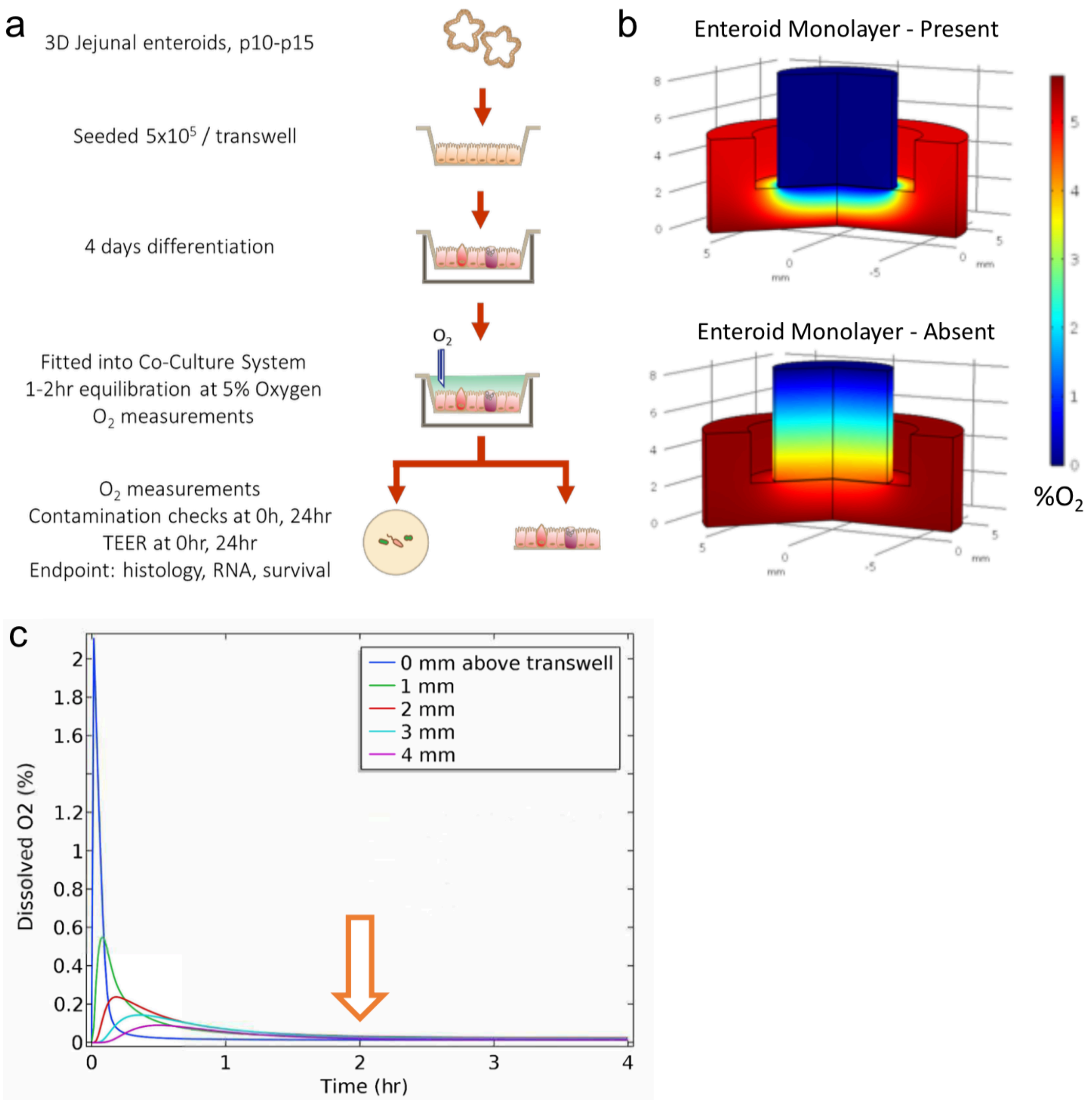


**Supplementary Figure 1. Enteroid-Anaerobe Co-Culture (EACC) experimental design and modeling.** (a) Schematic representation of the workflow for physiological hypoxia experiments. (b) Oxygen Consumption models show that enteroid monolayers scavenge basolateral oxygen (in red) and prevent leaks into the apical compartment (left panel). The right panel visualizes the gradual dissipation of dissolved oxygen in the absence of an enteroid monolayer. (c) Given the oxygen consumption capacity of enteroids, calculated time to equilibrium (red arrow) is approximately 2 hours.

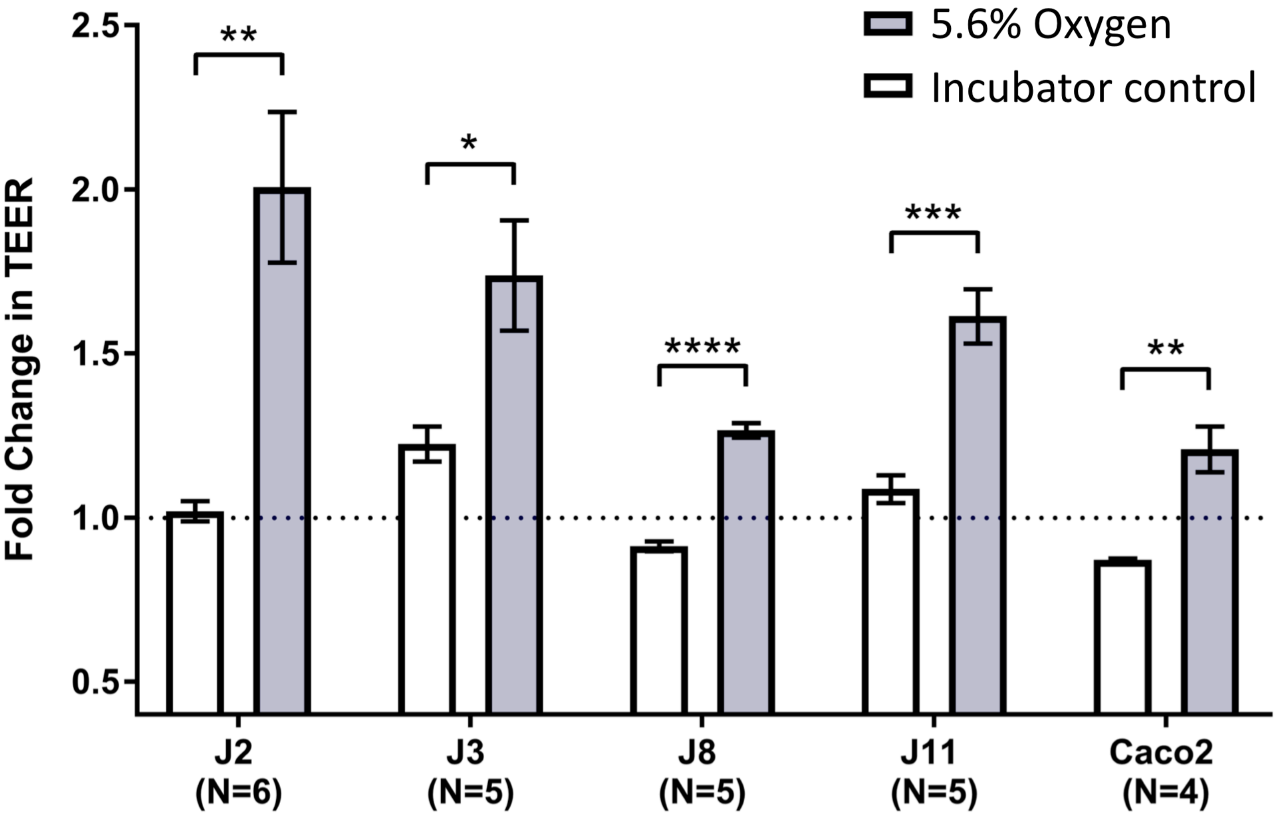


**Supplementary Figure 2. Enteroid response to physiological hypoxia.** The phenotype of increased barrier integrity in response to physiologic hypoxia is seen across 5 cell lines. Error bars reflect standard error; significance determined via unpaired, two-tailed t-test with a Benjamini & Hochberg FDR correction to obtain p-values (* $p<0.05$, ** $p<0.01$, ***$p<0.001$, **** $p<0.0001$).

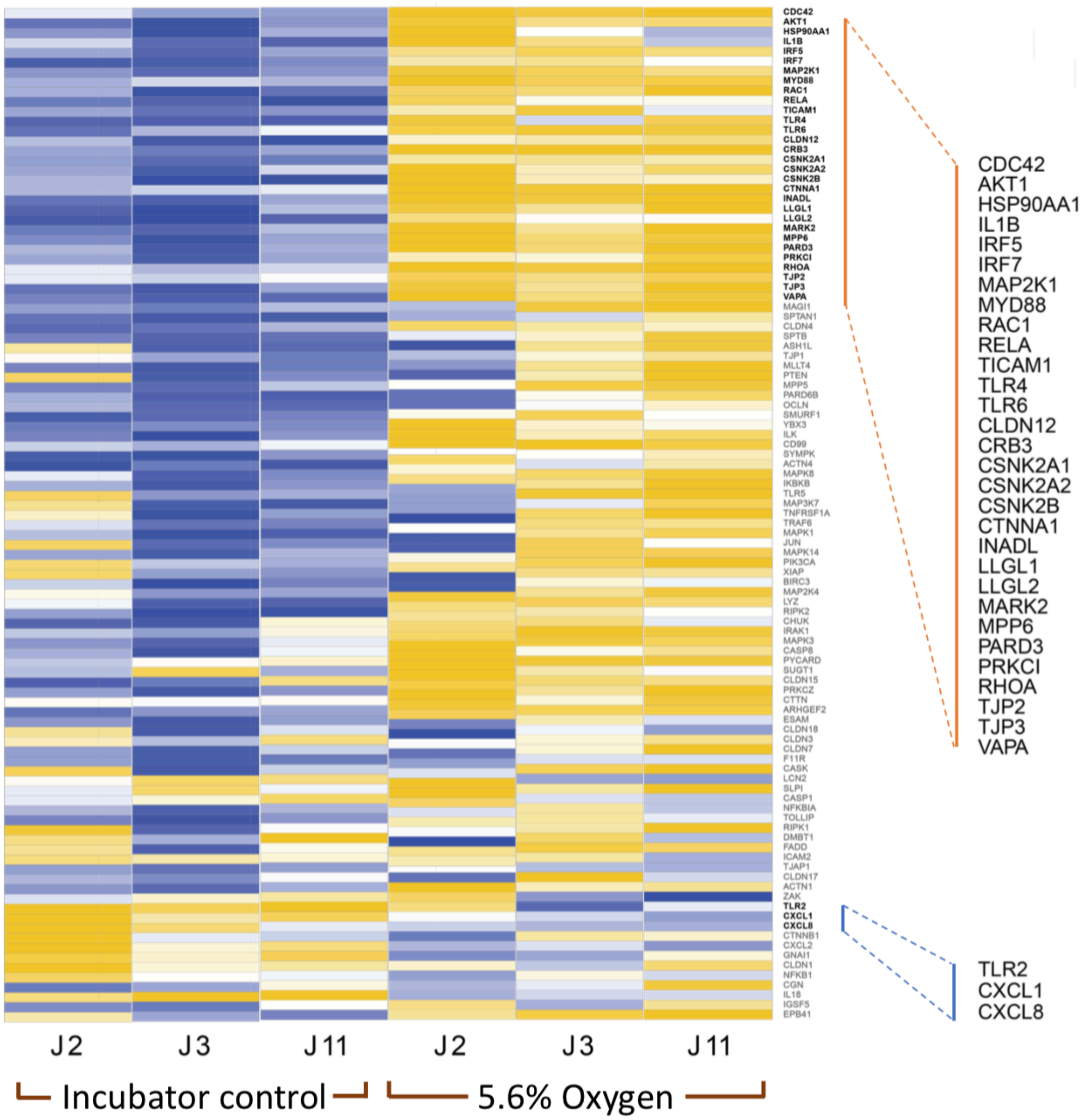


**Supplementary Figure 3. Exposure to physiological hypoxia for 24 hours results in a significant upregulation of epithelial integrity and innate immune response genes compared to standard incubator conditions.** Genes consistently upregulated >2 fold across all three jejunal lines in response to hypoxia in orange inset. Genes consistently downregulated >2 fold across all three jejunal lines in blue inset.

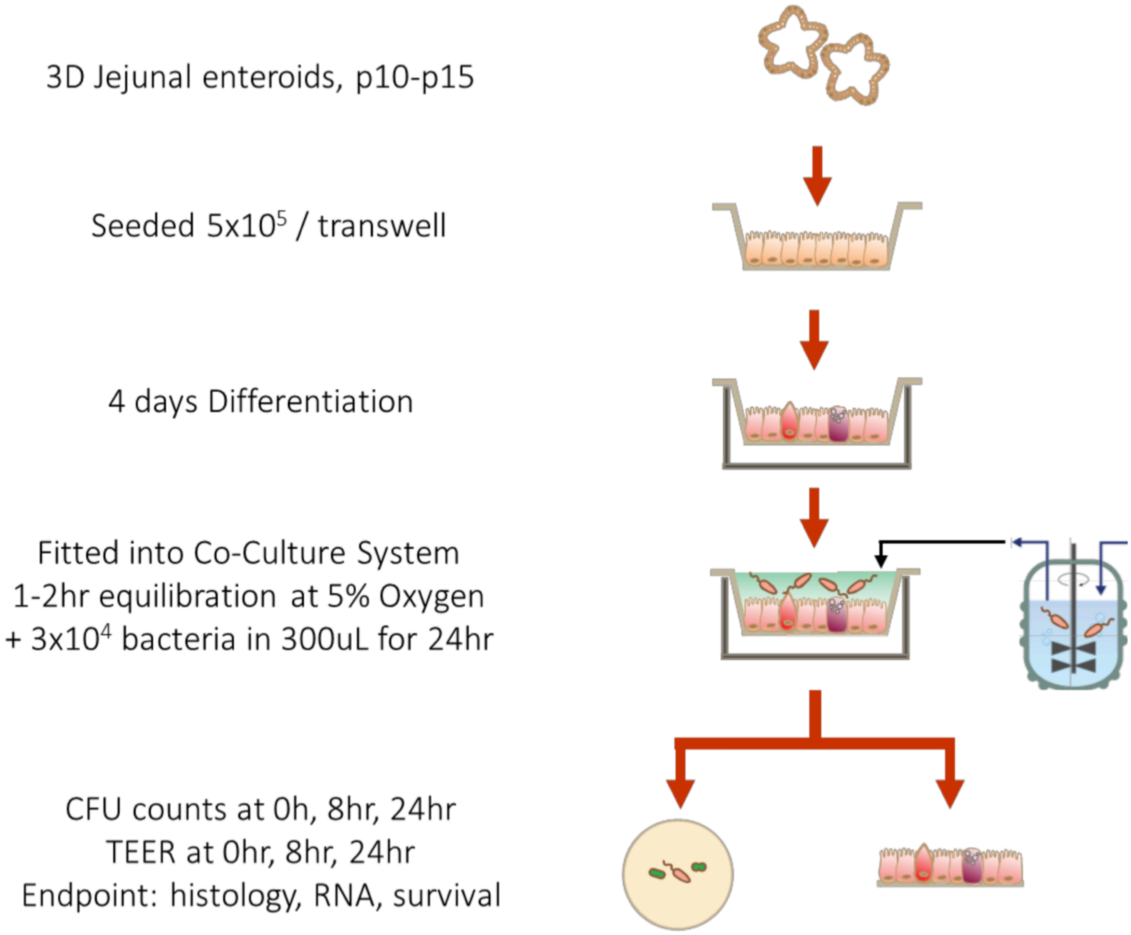


**Supplementary Figure 4. Schematic representation of the workflow for bacterial enteroid co-culture under hypoxia.**

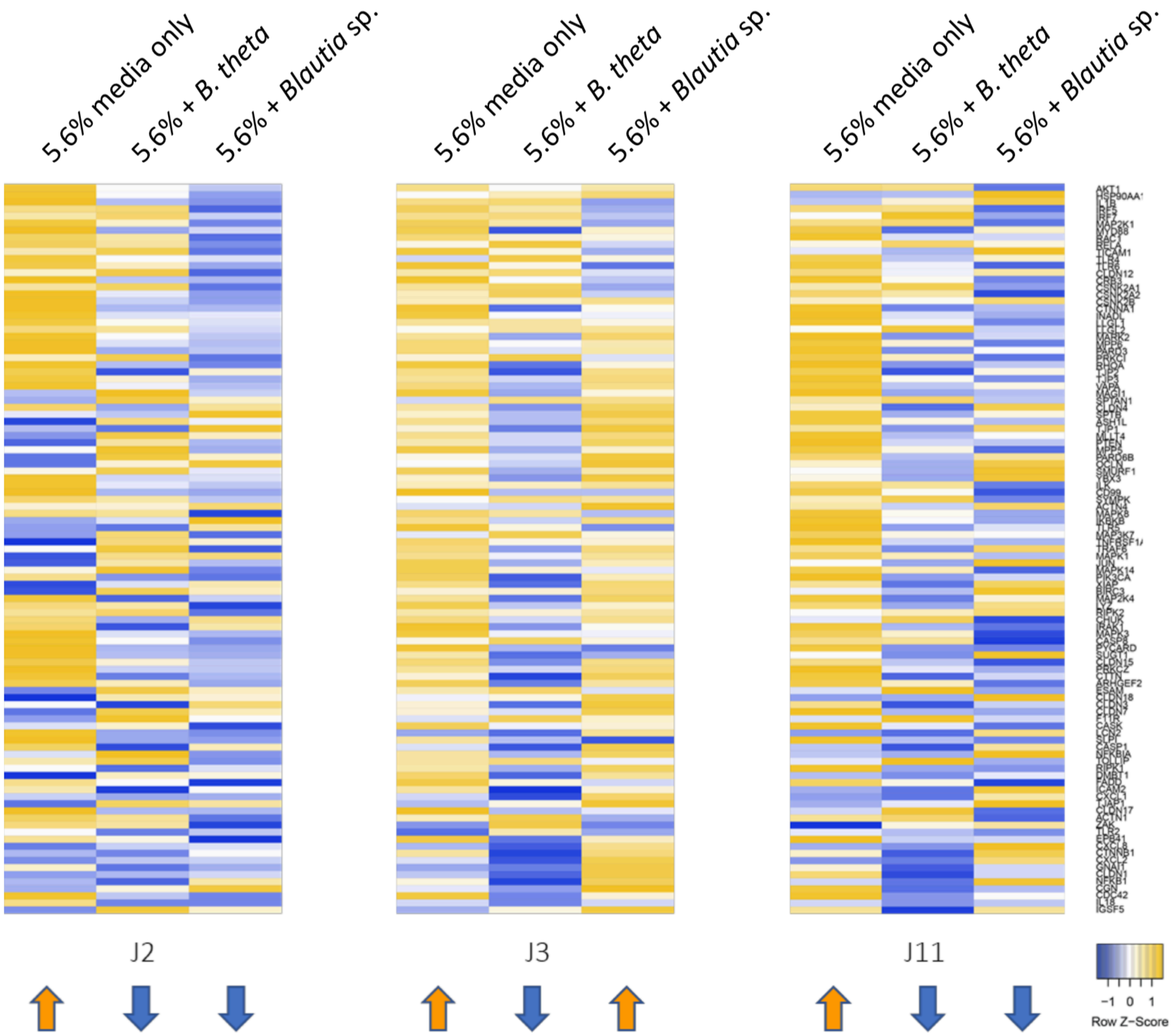


**Supplementary Figure 5. Patient-specific response to bacterial co-culture.** All three patient-derived lines have reduced expression of anti-microbial response and barrier integrity genes in response to *B. thetaiotaomicron (B. theta*) co-culture (relative to 5.6% hypoxia). However, these genes remain upregulated during *Blautia* co-culture in the J3 lineage, contrary to the J2 and J11 lines. General upregulation indicated by orange arrows; general downregulation indicated by blue arrows.

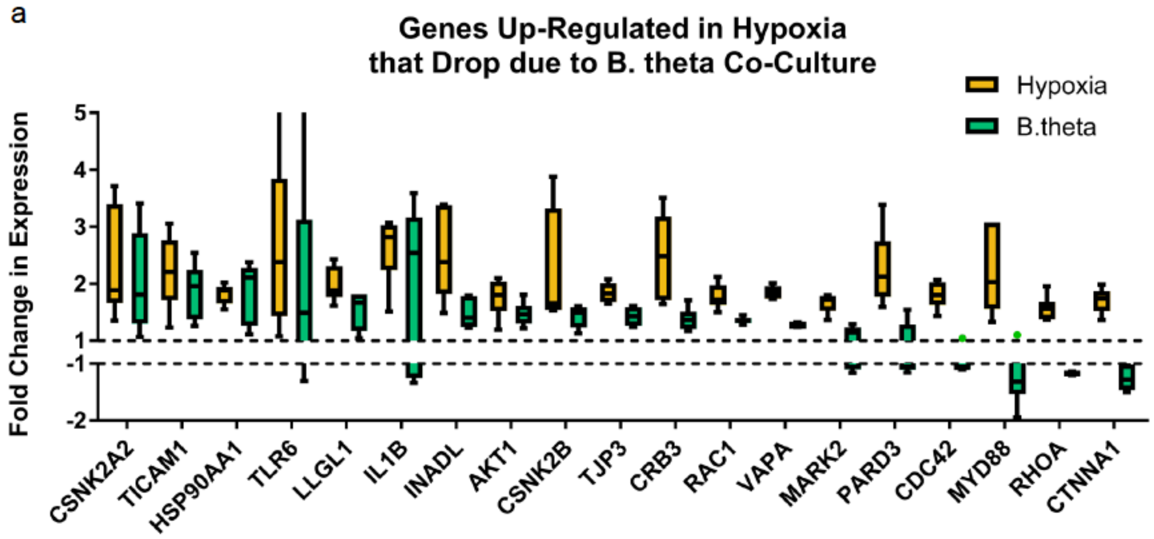


**Supplementary Figure 6. Enteroid response to *B. thetaiotaomicron* co-culture under physiological hypoxia.** Many of the genes that were upregulated during physiological hypoxia are mitigated following exposure to *B. thetaiotaomicron (B. theta*) co-culture.
